# HDL-AuNPs-BMS nanoparticle conjugates as molecularly targeted therapy for leukemia

**DOI:** 10.1101/250985

**Authors:** Na Shen, Fei Yan, Jiuxia Pang, Zhe Gao, Aref Al-Kali, Christy L. Haynes, Mark R. Litzow, Shujun Liu

## Abstract

In previous work, gold nanoparticles (AuNPs) with adsorbed high-density lipoprotein (HDL) nanoparticles have been utilized to deliver oligonucleotides, yet HDL-AuNPs functionalized with small molecule inhibitors have not been systematically explored. Here, we report an AuNP-based therapeutic system (HDL-AuNPs-BMS) for acute myeloid leukemia (AML) by delivering BMS309403 (BMS), a small molecule that selectively inhibits AML-promoting factor fatty acid binding protein 4 (FABP4). HDL-AuNPs-BMS are synthesized using a gold nanoparticle as template to control conjugate size and ensure a spherical shape to engineer HDL-like nanoparticle containing BMS. The zeta potential and size of the HDL-AuNPs obtained from transmission electron microscopy (TEM) show that the nanoparticles are electrostatically stable and 25 nm in diameter. Functionally, compared to free drug, HDL-AuNPs-BMS conjugates are more readily internalized by AML cells and have more pronounced effect on downregulation of DNA methyltransferase 1 (DNMT1), reduction of global DNA methylation, and restoration of epigenetically-silenced tumor suppressor *p15*^INK4b^ coupled with AML growth arrest. Importantly, systemic administration of HDL-AuNPs-BMS conjugates into AML-bearing mice inhibits DNMT1-dependent DNA methylation, induces AML cell differentiation and diminishes AML disease progression without obvious side effects. In summary, these data, for the first time, demonstrate HDL-AuNPs as an effective delivery platform with great potential to attach distinct inhibitors, and HDL-AuNPs-BMS conjugates as a promising therapeutic platform to treat leukemia.

## 1. INTRODUCTION

Gold nanoparticles (AuNPs) have emerged as attractive candidates for cancer therapeutic^1^ and diagnostic^2–3^ agents as well as drug delivery vehicles,^3–4^ because AuNPs are biocompatible, non-toxic and highly tolerable^5–6^. The subcellular size and good biocompatibility allow the circulating AuNPs to preferentially accumulate at tumor site,^7^ thereby selectively delivering therapeutic agents to cancer cells. They can be easily synthesized in various shapes, including spherical, rod-like and core–shell,^8^ with sizes ranging from 1 to 200 nm and with the spherical shape as the most useful form. AuNPs provide a highly multifunctional platform, which enables particle surfaces to be functionalized with various biological molecules.^9–11^ For instance, hydrophobic drugs can be loaded onto AuNPs through non-covalent interactions without structural modifications for drug release.

High density lipoproteins (HDLs) are “good” cholesterol^12^ that has been shown to be protective against the development of atherosclerosis and resultant illnesses.^13–14^ HDLs stimulate the growth of many types of cells, including cancer cells. However, the roles of HDLs in cancer are controversial. These investigations suggest that high levels of HDLs increase the risk of breast cancer, due to the high expression of the HDL receptor, scavenger receptor type BI (SRB1), on breast cancer cells.^15–16^ Notably, SRB1 is usually not expressed in the majority of normal human cells.^17^ However, the others have found that high HDL levels may protect against colon cancer incidence,^18^ but low HDL-C is associated with increased postmenopausal breast cancer risk.^19^ Further, HDLs are dynamic natural nanoparticles that transport cholesterol throughout the body and also other lipids, proteins and oligonucleotides.^12^ Indeed, HDLs transport and deliver endogenous microRNAs (miRs) in plasma to recipient cells with functional targeting capabilities.^20^ Owing to such promise as a delivery vehicle, mimicking HDL-NPs have been synthesized and utilized to deliver DNA, siRNA or small molecules *in vitro* and *in vivo*.^21–23^

While AuNPs offer a template to precisely control conjugate size, they don’t have a specific marker to be targeted on the surface of cancer cells. As a delivery vehicle, HDL-NPs could bind to SRB1, thereby selectively killing cancer cells by the loaded drug. Thus, the combination of AuNPs with HDL could generate cancer cell-specific NPs with controllable size and spherical shape. By using an AuNP core, Thaxton *et al* reported the synthesis of the first biomimetic HDL (HDL-AuNPs), with cholesterol binding capabilities, for therapeutic application.^24^ Other investigators showed that HDL-AuNP platform delivers chol-DNA antisense to repress *miR-210* expression in prostate cancer cells,^25^ and HDL-AuNPs conjugates can manipulate the efflux of cholesterol in lymphoma cells that leads to cell apoptosis and the selective inhibition of B-cell lymphoma growth.^26^ However, the utilization of HDL-AuNPs to deliver small-molecule drugs for leukemia therapy has not been initiated.

Leukemia is a heterogeneous blood cancer with diverse clinical and molecular features. Although a number of genetic and epigenetic abnormalities involved in leukemia pathogenesis have been identified and serve as potential therapeutic targets, outcomes are still dismal and relapse rates remain high in majority of leukemia patients. Fatty acid binding protein 4 (FABP4) is a key adipokine induced during adipocyte differentiation, and a fatty acid chaperone delivering fatty acids (FAs) to the nucleus. Our recent findings^27^ demonstrated that FABP4 upregulation in host and in leukemia cells facilitates leukemia expansion and invasion in both a cell-autonomous and cell-non-autonomous manner. Mechanistically, this process is mediated by FABP4-enhanced DNA methyltransferease 1 (DNMT1) expression and DNA methylation in AML cells. These findings highlight FABP4 as a novel therapeutic target for leukemia. BMS-309403 (BMS), a potent and selective FABP4 inhibitor, interacts with the fatty-acid binding pocket within the interior of FABP4 to block binding of fatty acids^28^. Targeting FABP4 by BMS for management of metabolic diseases has been investigated, but it remains unexplored whether and by which mechanism BMS has anti-leukemia properties. In current study, we synthesized and characterized HDL-AuNPs loaded with BMS (HDL-AuNPs-BMS). We tested the DNA hypomethylating activities and the anti-leukemia growth of HDL-AuNPs-BMS *in vitro* and *in vivo*. We showed that treatment with HDL-AuNPs-BMS blocks leukemia cell growth *in vitro* and induces leukemia regression *in vivo*, which occurs through a reduction of DNMT1-dependent DNA methylation. These findings suggest that the HDL-AuNPs are viable platforms to deliver small molecule into leukemia cells and the HDL-AuNPs-BMS has great potential for treatment of leukemia patients characterized by frequent epigenetic aberrations.

## 2. MATERIALS AND METHODS

### 2.1. Synthesis of Au-HDL-NP

The HDL-AuNPs citrate-stabilized colloidal gold nanoparticles (AuNPs) were synthesized similarly to published Ji’s protocol with slight modifications.^29^ The AuNPs were then washed and collected by centrifuging at 8000 rpm for 15 min. The ultrapure (18.2 MΩ) water was used for synthesis of the HDL-AuNPs and all other experiments.

Sulfhydryl groups were added to Apo AI using Traut’s reagent to modify the protein for nanoparticle attachment. Traut’s reagent reacts with the primary amines that reside on lysine residues to generate amino acids with sulfhydryl groups. Apo AI was reconstituted in a small amount of PBS containing 4 mM EDTA at pH 8.0. A 20-fold molar excess of Traut’s reagent was added and allowed to incubate with the protein for 1 hour at room temperature (RT). The modified Apo AI (Apo AI-SH) was then purified with a Zeba column.

For the HDL-AuNPs containing Apo AI-SH, the protein was added to the Au colloid in a 135-fold molar excess. After incubation with Apo AI-SH (10 hours at room temperature), the phospholipids were added. The phospholipids were dissolved in ethanol, and fresh solutions of 1 mM were used for each synthesis. The AuNPs containing Apo AI-SH were diluted 50% with ethanol. Two phospholipids, 1,2-dipalmitoyl-sn-glycero-3-phosphoethanolamine-N-[3-(2-phyridyldithio) propionate] (PDP PE) and 1,2-dipalmitoyl-sn-glycero-3-phosphocholine (DPPC), were both added in 15000-fold molar excess. The solutions were incubated (10 hours, room temperature) and subsequently purified by diafiltration using a KrosFlo Research II tangential flow filtration system (Spectrum Laboratories) fitted with a 50 kDa modified polyethersulfone (mPES) module. For all purifications, the buffer was exchanged at least seven times to remove unreacted Apo AI-SH and phospholipids. An aqueous solution of 50% ethanol was used at the beginning of the purification and changed to 25% ethanol and, ultimately, to water to aid in phospholipid removal.

### 2.2. Transmission electron microscopy

About 3 µl aliquot of concentrated Au colloid or Au-lipid sample was added to glow discharged TEM carbon grid. The grid was either blotted dry or stained with 2% uranyl acetate. Images were recorded at a nominal magnification of 115,000 × using a FEI Tecnai 300-kV F30 field emission gun transmission and viewed through a FEI electron microscope; pictures were taken using a CCD camera.

### 2.3. Determination of BMS in the HDL-AuNPs nanocarrier

The BMS-loading content, defined as the weight percentage of BMS in the nanocarrier, was quantified with a UV-vis 2550 spectrophotometer (Shimadzu, Japan). First, solutions of the BMS-loaded nanocarriers were centrifuged, and the supernatant was collected. After that, the free BMS was added to equal volume of ethanol. Then, absorbance at 260 nm was measured to determine the drug content in the solution from a previously established calibration curve. The unloaded content of BMS was calculated and the encapsulation efficiency (EE) was obtained according to the following equation:

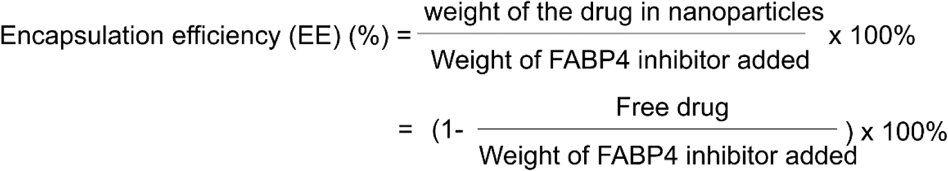

### 2.4. Cell culture

Cell lines Kasumi-1, MV4-11, K562, C1498, and Hep3B were obtained from American Type Culture Collection (Manassas, VA, USA). All cell lines were cultured in DMEM (C1498) or RPMI1640 (others) medium supplemented with 20% (Kasumi-1) or 10% (others) fetal bovine serum at 37°C under 5% CO_2_. FABP4 inhibitor (BMS309403) was purchased from Calbiochem and dissolved in ethanol for pre-clinical test and in DMSO for cell culture experiments.

### 2.5. Methylcellulose colony-forming and MTS assays

Colony-forming assays were performed in MethoCult^®^ mixture (Stem Cell Technologies Inc), and MTS assays were conducted using CellTiter 96^®^ AQueous One Solution Cell Proliferation Assay (Promega), according to the manufacturer’s instructions. The number of colonies were scored in 7–14 days.

### 2.6. Analysis of apoptosis

Apoptosis was measured using the Annexin V-FITC apoptosis detection Kit (BD Biosciences, Pharmingen) as previously described^30–31^. Briefly, MV4-11, C1498 and bone marrow (BM) cells were treated with free BMS in ethanol, empty NPs or HDL-AuNPs-BMS for 48 hours, stained with Annexin V-FITC and propidium iodide and analyzed by flow cytometry using FACSCalibur (BD Biosciences). Data were analyzed using CellQuest 3.3 software (BD Biosciences).

### 2.7. Cytospin/Wrigth-Giemsa staining

At the designated time points, 0.1 × 10^6^ of MV4-11, C1498 and BM cells were harvested and placed in Shandon EZ Single Cytofunnel (Thermo Electron Corporation). Samples were centrifuged at 1,000 rpm for 8 min. The slides were air-dried and stained with Hema-3 Kit (Fisher Scientific) following the protocol described previously^27, 30, 32^.

### 2.8. Western blot

After different treatment, the whole cellular lysates were prepared by harvesting the cells in 1 × cell lysis buffer and the Western blot was performed as previously described^27, 30, 33–34^. Equivalent gel loading was confirmed by normalizing to β-actin levels. The antibodies used are: β-actin (Santa Cruz Biotechnology), DNMT1 (New England Biolabs, Ipswich, MA).

### 2.9. Dot blotting

The genomic DNA was purified using DNA Blood/Tissue Kit (QIAGEN) and Dot blotting was performed as previously described^27, 30, 32, 34^. Around 2 μg of DNA was denatured in 1 × buffer (0.4M NaOH, 10mM EDTA) at 100°C for 10 minutes and spotted onto the nylon by heating at 80°C for 2 hours. The DNA spotted membrane was blocked with 5% nonfat milk for 1 hour and probed with mouse anti-5mC (1:2500, Active Motif, CA) and the signal was detected by HRP-conjugated secondary antibody and enhanced chemiluminescence.

### 2.10. RNA isolation, cDNA preparation and quantitative PCR

According to the manufacturer’s instructions, RNA was isolated using miRNAeasy Kit (QIAGEN) and cDNA synthesis was performed using SuperScript^®^ III First-Strand Synthesis System (Invitrogen). For *DNMT1* expression, qRT-PCR was carried out by TaqMan technology (Applied Biosystems, Foster City, CA). *18S* levels were analyzed for normalization. Expression of the target genes were measured using the ΔCT approach. Primers used are:

*p15*^INK4B^: forward 5’-CCAGATGAGGACAATGAG-3’;

*p15*^INK4B^: reverse 5’-AGCAAGACAACCATAATCA-3’;

*h18S*: forward 5’-ACAGGATTGACAGATTGA-3’;

*h18S*: reverse 5’-TATCGGAATTAACCAGACA-3’;

*m18S*: forward 5’-ACAGGATTGACAGATTGA-3’;

*m18S*: reverse 5’-TATCGGAATTAAC CAGACA-3’.

### 2.11. Bisulfite sequencing

About 2 μg of total DNA was bisulfite-converted and purified using EpiTect Bisulfite Kit (Qiagen). The region (-4 to +247) within the *p15*^*INK4B*^ promoter was amplified by PCR using the following primers: forward 5’-GGTTGGTTTTTTATTTTGTTAGA G-3’; reverse 5’-ACCTAAACTCAACTTCATTACCCTC-3’. The PCR products were subcloned using the TA Cloning^®^ Kit (Invitrogen), and sequenced in Genewiz company^27, 30, 32, 35–36^.

### 2.12. Animal studies

All animal experiments were approved by the Institutional Animal Care and Use Committees of the University of Minnesota and were in accordance with the U.S. National Institutes of Health (NIH) Guide for Care and Use of Laboratory Animals. For the preclinical testing of the free FABP4 inhibitor BMS309403, the drug was prepared just prior to administration by dissolving it in ethanol to give a clear solution and diluting with PEG400 and saline (ratio 15:38:47). Six C57BL/6 mice (male, aged 4–6 weeks) were divided into two groups (n = 3), and C1498 cells (0.1 × 10^6^) were engrafted by tail-vein injection. When the white blood cell (WBC) count indicated the development of leukemia, the drug BMS309403 (free or HDL-AuNPs-BMS) was given via tail-vein in every three days: four times at 5 mg/kg and two times at 10 mg/kg. The same administration of vehicle was used for the negative control. For WBC counting, 2 μl of blood from mouse tail vein was mixed with 38 μl of Turk blood dilution fluid (Ricca Chemical), and the WBCs were counted under microscope. Spleen weight and number of metastatic nodules were determined at the end of each experiment and tissues were fixed in 10% formalin.

### 2.13. Hematoxylin and Eosin (H&E) staining

Tissues collected from our animal studies were fixed in 4% paraformaldehyde/PBS. The paraffin-embedded samples were cut to 5 μm thick and H&E staining were performed as previously described^27, 30, 32^. Samples were developed with 3, 3′-diaminobenzidine (Vector Laboratories), counterstained with hematoxylin, and mounted. Stained slides were viewed and photographed with a Leica microscope mounted with a high-resolution spot camera, which is interfaced with a computer loaded with Image-Pro Plus software.

### 2.14. GEO (Gene Expression Omnibus) analysis

AML GEO dataset (GSE12417, platform GPL570, n=163) were analyzed for the mRNA expression of SRB1, which was assessed by gene-expression arrays. The patients reported in this GEO are AML and cytogenetically normal. The detailed clinical characteristics of the patients were referred to the original report.^37^ These samples were normalized, managed and analyzed by GraphPad Prism 5 Software using Spearman correlation coefficients.

### 2.15. Statistical analysis

The Western blot, real time PCR, Dot blotting and organ weight data were analyzed using the Student’s t-Tests. All statistical analysis was carried out using GraphPad Prism 5.0. Differences were considered statistically significant at *P* < 0.05. All *P* values were two-sided.

## 3. RESULTS AND DISCUSSION

### 3.1. SRB1 is highly expressed in leukemia cells

In order to maintain a high level of growth, cancer cells (e.g., breast cancer) scavenge HDL particles by overexpressing its receptor, SRB1.^16^ It is reported that myeloblasts from the patients with AML have higher cholesterol uptake via HDL carriers,^38^ and disruption of cholesteryl ester formation inhibits leukemia cell growth.^39^ In the present study, we found that SRB1 is largely detectable in leukemia cells lines, Kasumi-1, MV4-11, and K562, when compared to the hepatic cell Hep3B (Figure 1a), which highly expresses SRB1.^16, 40–41^ Importantly, higher SRB1 expression is associated with a shorter survival time in AML patients (Figure 1b), supporting the pathogenic contribution of SRB1 to AML disease progression. However, it has been observed that the decreased levels of HDL-C were seen in plasma of leukemia patients,^39^ supporting the idea that HDL-C may negatively regulate leukemogenesis. In addition, the circulating levels of FABP4, a regulator of lipid metabolism, are highly upregulated in obese patients^42^ and are inversely correlated to HDL-C levels.^43^ In line with this, upregulation of host and cellular FABP4 accelerates AML growth *in vitro* and *in vivo* in both a cell-autonomous and cell-non-autonomous manner.^27^

**Figure 1.**
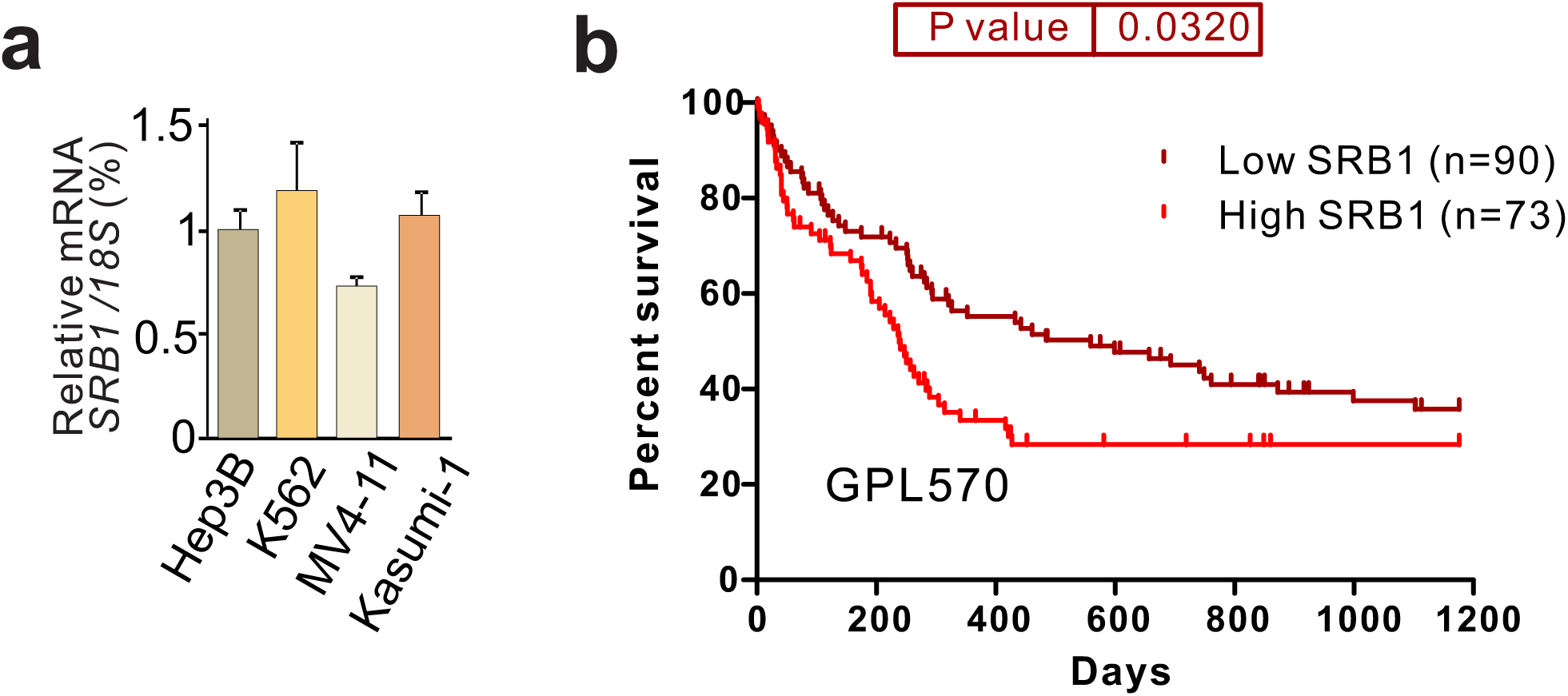
Upregulation of SCARB1 predicts worse outcomes. (a) qPCR measuring expression of SCARB1 in leukemia or hepatocellular carcinoma cells. (b) The analysis of the GEO dataset GSE12417, platform GPL570, containing 163 patients with AML. The patient overall survival (OS) was analyzed using Kaplan–Meier estimate.

### 3.2. Synthesis and characterization of HDL-AuNPs-BMS

BMS309403 (BMS), a biphenyl azole inhibitor, specifically binds and inactivates FABP4^28^. Although BMS is effective in curing certain metabolism disease^44^, its therapeutic potential in cancers, particularly in the context of nanoparticles, remains elusive. To achieve a nanoparticle-based HDL therapeutic platform, we employed gold nanoparticles (AuNPs) as a scaffold to synthesize HDL biomimics. These AuNPs can accurately control the size, shape and surface chemistry of spherical HDL (Figure 2a,b). The methods for AuNP synthesis have been modified from a previous report.^29^ The main surface components of natural HDL, like apolipoprotein AI (Apo AI) and phospholipids, were loaded onto the AuNP surface. HDL-AuNPs were characterized in terms of stability and diameter. HDL-AuNPs were stable in solution, because they were red in color and have a measured SPB of 520 nm in the ultraviolet visible spectrum (Figure 2c). TEM characterization showed that the sizes of AuNPs were 25 nm in diameter (Figure 2d), which was the suitable size to achieve enhanced permeability and retention (EPR) effect.^45^ Notably, all these constructs mimic the general physical and structural properties of natural HDL.

**Figure 2.**
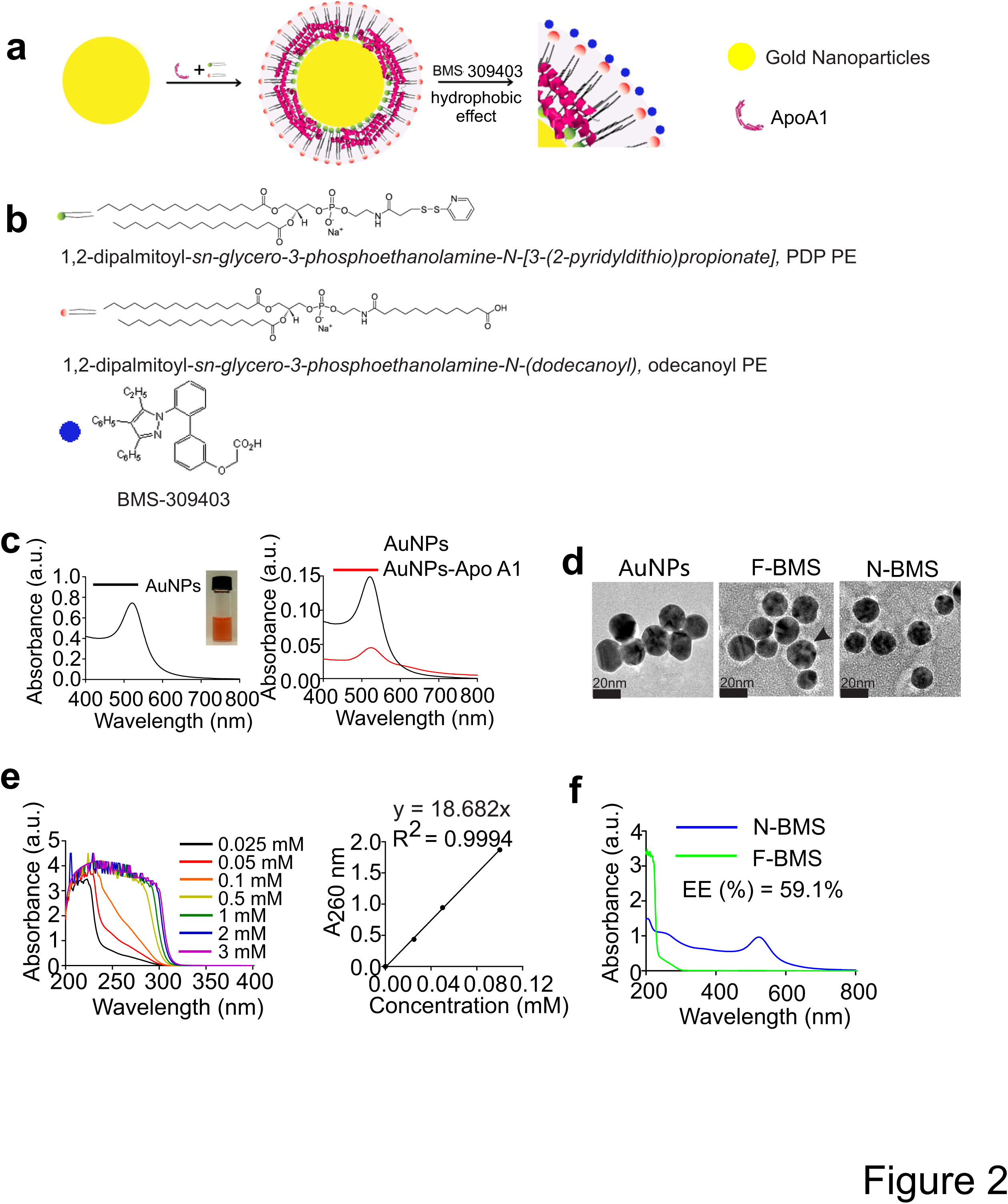
Synthesis and characterization of HDL-AuNPs-BMS. (a) Scheme of HDL-AuNPs-BMS synthesis. Colloidal AuNPs were mixed with apolipoprotein A-I (APOAI) followed by the addition of phospholipids to form biomimetic HDL-AuNPs. The HDL-AuNPs were purified and resuspended in water. (b) The structure of two phospholipids and BMS. (c) The UV-Vis spectra of AuNPs and Apo-A1 modified AuNPs. (d) Transmission electron micrographs showing the size of individual AuNPs, HDL-AuNPs and HDL-AuNPs-BMS. Arrow indicates the lipid layer. (e) Spectrometry and standard curve of BMS. (f) The UV-vis spectroscopy for the stability of the HDL-AuNPs-BMS in PBS, and free BMS. Note: Emp, F-BMS, N-BMS represents empty vehicle, free BMS, HDL-AuNPs-BMS, respectively.

Having fabricated HDL-AuNPs as described in scheme (see Figure 2a and 2b), next we sought to choose a small-molecule drug and construct HDL-AuNPs-drug conjugates. We selected the hydrophobic compound, BMS, because it is a selective inhibitor for FABP4^28^, whose deregulation plays a key role in leukemia pathogenesis^27^. Initially, BMS was incubated with HDL-AuNPs for 6 hours. The product HDL-AuNPs-BMS was purified and resuspended in PBS. The UV-vis spectroscopy verified the stability of the HDL-AuNPs and AuNPs in PBS. A surface plasmon band centered at ~520 nm, which is consistent with disperse AuNPs (Figure 2c). TEM data showed the difference of thickness between the monolayer and bilayer lipid functionalization (Figure 2d). The measurement of dynamic light scattering (DLS) revealed little size increase after BMS addition (25 ± 4 nm). The UV-Vis spectrum of BMS with different concentration was shown, and the standard curve between concentration and absorbance at 260 nm was obtained (Figure 2e). Moreover, the absorption band of HDL-AuNPs-BMS displayed a slight lift at 200-300 nm, which was the contribution of free BMS (Figure 2f), collectively, supporting the successful synthesis of the HDL-AuNPs-BMS conjugates.

### 3.3. Treatment with HDL-AuNPs-BMS induces growth arrest of leukemia cells

To assess the ability of HDL-AuNPs-BMS to suppress leukemia cell growth, AML cell lines, MV4-11 and C1498, and bone marrow (BM) cells from leukemia-bearing mice were treated with free BMS in ethanol, bare AuNPs or HDL-AuNPs-BMS. Colony-forming assays showed that HDL-AuNPs-BMS induced the most robust decrease of colony number compared to free BMS and bare NPs (Figure 3a). The MTS analysis further demonstrated the most significant impairment of cell proliferation by HDL-AuNPs-BMS in comparison with their respective controls (Figure 3b). Because leukemia cells are developed from a blockade of differentiation at a distinct stage in cellular maturation, one major goal of chemotherapy is to force leukemia cells to undergo terminal differentiation.^46^ To this end, we employed Wright-Giemsa to stain the above treated leukemia cells^27, 30^. In response to HDL-AuNPs-BMS, MV4-11, C1498 and BM cells displayed the highest potential of cell differentiation, as indicated by more post-mitotic cells containing metamyelocytes, bands and segmented neutrophils (Figure 3c). However, flow cytometry analysis showed that neither HDL-AuNPs-BMS nor free BMS had obvious effects on cell apoptosis in MV4-11, C1498 and BM cells (Figure 3d). These results support that HDL-AuNPs-BMS conjugates have higher efficacy in repressing leukemia cell growth *in vitro* through an induction of cell differentiation.

**Figure 3.**
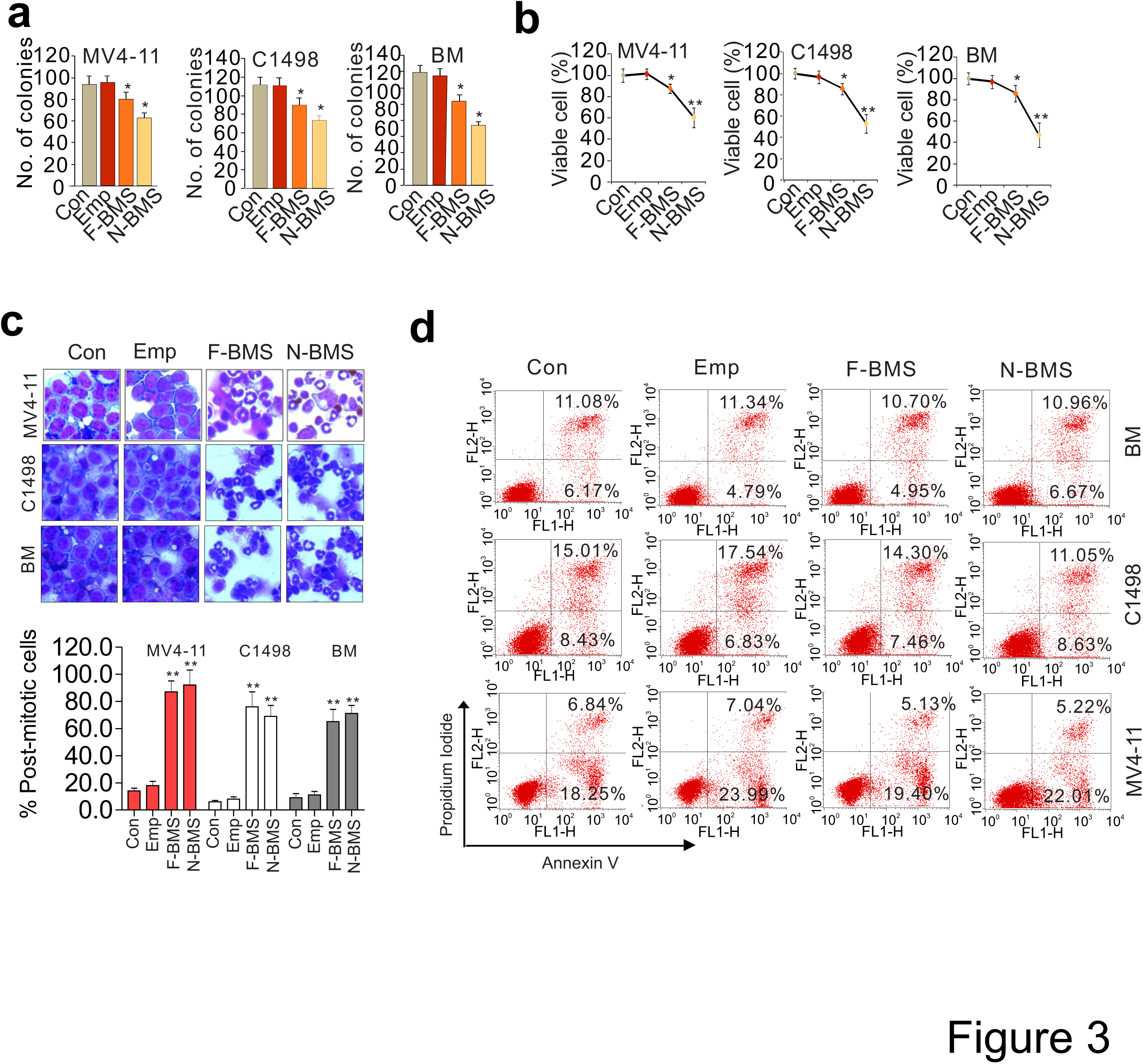
Inhibitory effect of HDL-AuNPs-BMS on leukemia cell growth *in vitro*. (a) Colony-forming assays in MV4-11, C1498 or mouse BM cells treated with empty NPs or 10 µM HDL-AuNPs-BMS and free BMS for 48 hours. Graph indicates the colony number from three independent experiments. Data are mean ± SD, \**P* < 0.05. (b) MTS cell proliferation assays in MV4-11, C1498 or mouse BM cells treated with empty NPs or 10 µM HDL-AuNPs-BMS and free BMS for 72 hours. Data are mean ± SD, \**P* < 0.05, \*\**P* < 0.01. (c) Visual analysis of Wright-Giemsa-stained cytospins of MV4-11 and C1498 cells treated with empty NPs or 10 µM HDL-AuNPs-BMS and free BMS for 96 hours (original magnification × 400). Graphs show the count of post-mitotic cells. Data are mean ± SD, \*\**P* < 0.01. (d) Flow cytometry analysis in MV4-11 and C1498 cells treated with empty NPs or 10 µM HDL-AuNPs-BMS and free BMS for 72 hours. The experiments were performed three times independently. Note: Emp, F-BMS, N-BMS represents empty vehicle, free BMS, HDL-AuNPs-BMS, respectively.

### 3.4. Exposure of leukemia cells to HDL-AuNPs-BMS reduces DNMT1-dependent DNA methylation

To elucidate the mechanisms underlying the anti-leukemia properties of HDL-AuNPs-BMS, we subjected the protein lysates and RNA from the above treated cells to Western blot or qPCR.^30, 32^ In agreement with the role of FABP4 in DNMT1 regulation,^27^ exposure of MV4-11, C1498 and BM cells to HDL-AuNPs-BMS resulted in DNMT1 downregulation to the most at both RNA (Figure 4a) and protein levels (Figure 4b), compared to free BMS and vehicle NPs. We then prepared genomic DNA and utilized Dotblot^30, 35^ to measure changes in DNA methylation. DNMT1 downregulation by HDL-AuNPs-BMS was accompanied by substantial decrease of DNA methylation in these tested cells (Figure 4c). Notably, *p15*^*INK4B*^ is a tumor suppressor gene that is epigenetically silenced in leukemia,^47–48^ we reasoned that HDL-AuNPs-BMS may reactivate *p15*^*INK4B*^ through promoter DNA hypomethylation. Indeed, data demonstrated that *p15*^*INK4B*^ expression was increased (2.06 ± 0.11, 1.63 ± 0.105, 1.86 ± 0.088 folds, respectively) in MV4-11, C1498 and BM cells treated with HDL-AuNPs-BMS for 48 hours, which was superior to that of free BMS or vehicle NPs (Figure 4d). When the genomic DNA was bisulfite-converted and PCR-amplified, the sequencing of 10 clones disclosed 12% reduction of DNA methylation in HDL-AuNPs-BMS group, compared to those in vehicle NPs (Figure 4e).

**Figure 4.**
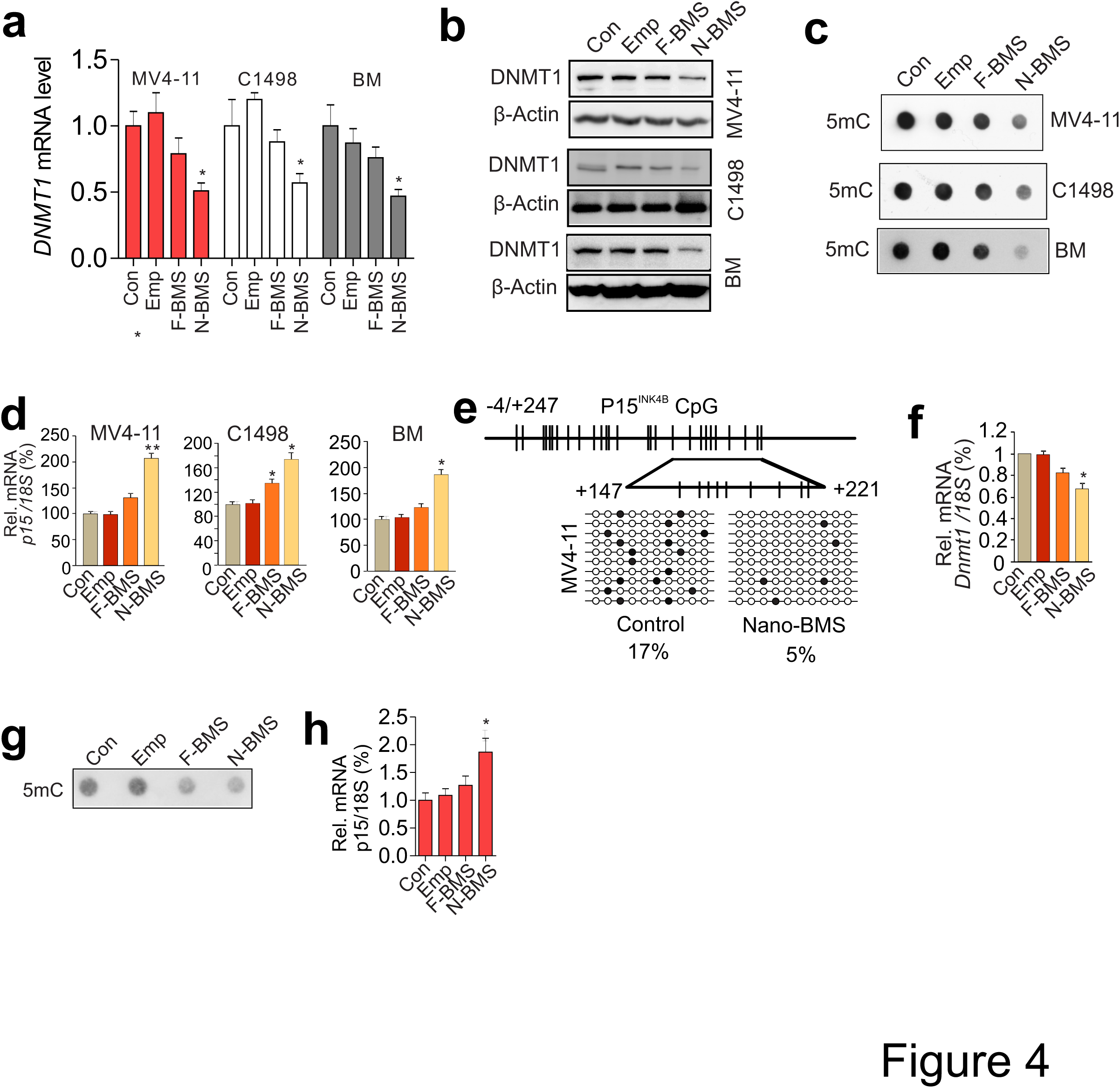
HDL-AuNPs-BMS displays DNA hypomethylating activities in leukemia cell. (a,b)qPCR (a) or Western blot (b) analysis for the expression of DNMT1 in MV4-11, C1498 or mouse BM cells treated with empty NPs or 10 µM HDL-AuNPs-BMS and free BMS for 48 hours. (c) Dotblot analysis using 5mC antibody for the change of DNA methylation in genomic DNA from MV4-11, C1498 or mouse BM cells treated with empty NPs or 10 µM HDL-AuNPs-BMS and free BMS 48 hours. (d) qPCR analysis of *p15*^*INK4B*^ expression in MV4-11, C1498 or mouse BM cells treated with empty NPs or 10 µM HDL-AuNPs-BMS and free BMS for 48 hours. Data are mean ± SD, \**P* < 0.05, \*\**P* < 0.01. (e) Bisulfite sequencing of *p15*^*INK4B*^ promoter in MV4-11 cells treated with empty NPs or 10 µM HDL-AuNPs-BMS and free BMS for 48 hours. CpG locations are indicated as vertical bars in the promoter and first exon of *p15*^*INK4B*^ (upper). Arrows indicate the bisulfite sequencing region (transcription start site +147 to +221). Lower: Open circles indicate unmethylated CpG sites, solid circles indicate methylated CpG sites. Results of 10 clones are presented. (e-g) Primary cells from AML patients (n = 3) were treated with empty NPs or 10 µM HDL-AuNPs-BMS and free BMS for 48 hours, and subjected to qPCR (f and h) for gene expression and Dotblot (g) for genomic DNA methylation. Note: Emp, F-BMS, N-BMS represents empty vehicle, free BMS, HDL-AuNPs-BMS, respectively.

To demonstrate the clinical implication of these findings, we treated primary cells from human AML patients (n = 3) with free or HDL-AuNPs-BMS. Consistent with the results from AML cell lines and mouse BM cells, the HDL-AuNPs-BMS induced a more pronounced reduction of DNMT1 expression (Figure 4f) and global DNA methylation (Figure 4g), but a greater increase of *p15*^*INK4B*^ as compared to free BMS (Figure 4h). These results provide compelling evidence of the efficacy of HDL-AuNPs-BMS in modifying the epigenetic profile in AML cells.

### 3.5. HDL-AuNPs-BMS therapy induces leukemia regression through DNA hypomethylation

To exploit the anti-leukemia activity of the HDL-AuNPs-BMS conjugates *in vivo*, we intravenously injected C1498 (0.1 × 10^6^) cells, a murine AML cell line that is syngeneic to C57BL/6 background,^27, 30, 34^ into C57BL/6 mice (n = 3) to mimic leukemic disease. When the WBC counts indicated the development of leukemia, the leukemia-bearing mice were given four doses of 5 mg/kg followed by two doses of 10 mg/kg HDL-AuNPs-BMS or free BMS (twice/week) in formulation of ethanol-PEG400-saline (ratio: 15:38:47).^30^ The age-matched leukemia-bearing mice (n = 3) injected with vehicle only were used as a negative control. When compared to vehicle-or free BMS-treated mice, the mice receiving HDL-AuNPs-BMS had the lowest number of WBCs (Figure 5a) and the smallest spleen (634 ± 33; 411 ± 21; 279 ± 19; Figure 5b). The quantification of metastatic foci revealed that the mice treated with HDL-AuNPs-BMS had the least nodules and the smallest lesion areas in lungs (19 ± 4; 7 ± 3; 3 ± 2) and livers (20 ± 5; 10 ± 3; 6 ± 2; Figure 5c and 5d), an indicator of suppressed lung and liver metastatic growth. H&E-stained sections of spleens, lungs and livers from HDL-AuNPs-BMS-treated mice showed no obvious infiltration of leukemic blasts, which were readily detectable in BM and organs of free- or vehicle-treated subjects (Figure 5e and 5f). Giemsa–stained BM showed that HDL-AuNPs-BMS therapy efficiently rescued granulocytic differentiation of myeloid cells. In fact, the majority of BM cells from HDL-AuNPs-BMS-treated mice were post-mitotic cells containing metamyelocytes, bands and segmented neutrophils (27 ± 5%; 57 ± 12%; 71 ± 19%; Figure 5g). Mechanistic investigation revealed that HDL-AuNPs-BMS administration resulted in more downregulation of DNMT1 expression (Figure 5h) and reduction of DNA methylation (Figure 5i), which was accompanied by a greater upregulation of methylation-silenced *p15*^*INK4B*^ gene (Figure 5j). Collectively, our results support the idea that the better antileukemic effects of HDL-AuNPs-BMS *in vivo* result from its greater capability to induce DNA hypomethylation in leukemia cells.

**Figure 5.**
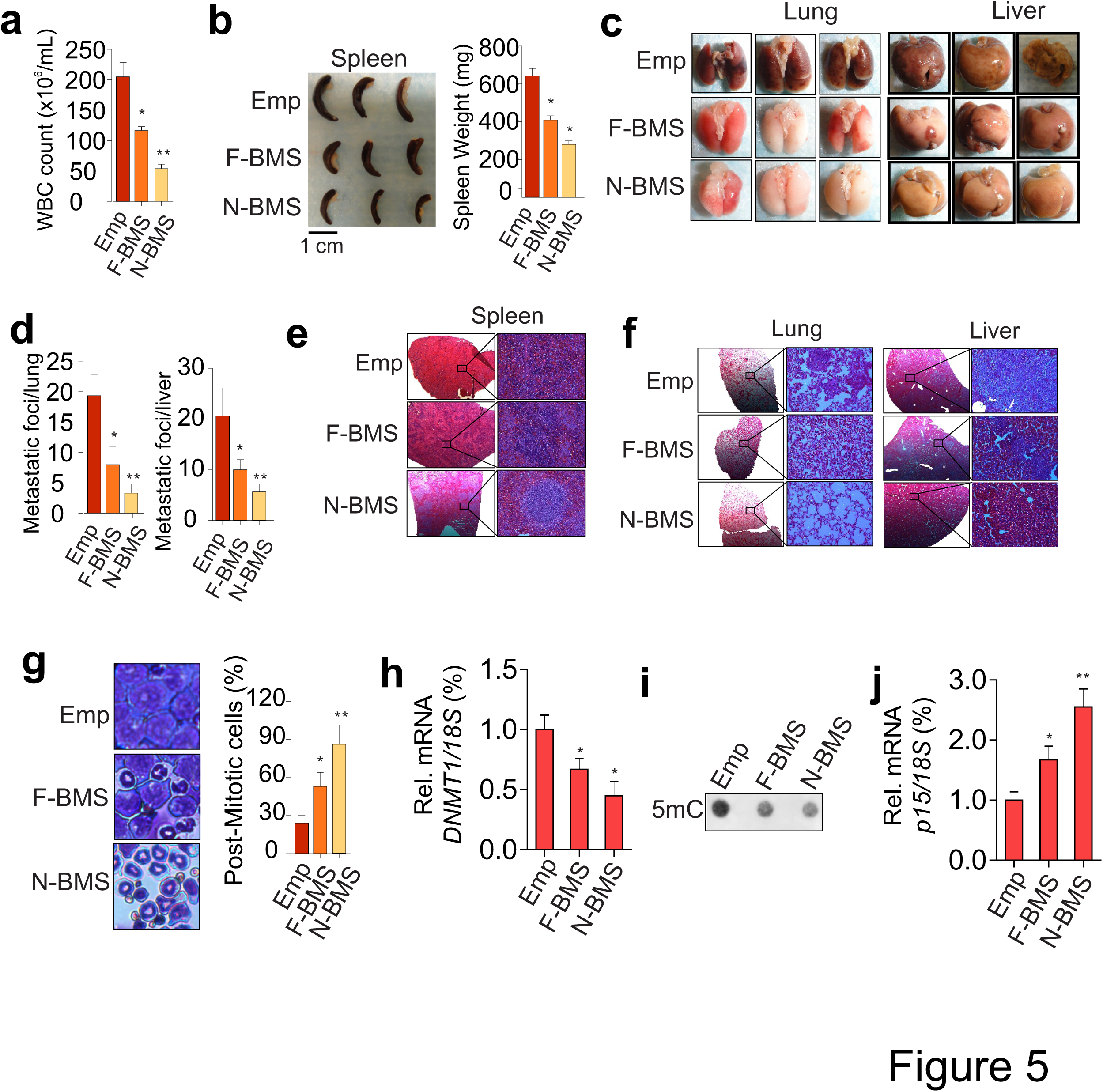
HDL-AuNPs-BMS administration *in vivo* attenuates leukemogenesis. (a) C1498 cells(0.1 × 10^6^) were intravenously injected into 4–6 weeks old C57BL/6 mice. When the mice showed leukemic disease, free BMS or HDL-AuNPs-BMS was initiated at the dose of 5 mg/kg. A twice-weekly treatment with 5 mg/kg/mouse for 2 weeks, which was continued at the dose of 10 mg/kg/mouse for 1 week while monitoring for any clinical signs of toxicity or leukemia-related distress. (b) Representative visual analysis (left) and calculated weight (right) of spleens from leukemia mice receiving free BMS, HDL-AuNPs-BMS or vehicle. Scale bars, 5.0 mm. (c) External views of lung and liver of untreated and drug-treated leukemia mice. (d) Graph is the quantification of tumor nodules growing in lung or liver. (e,f) H&E staining of tissue sections from lung or liver of leukemia mice receiving free BMS, HDL-AuNPs-BMS or vehicle. Scale bars, 100 μm. (g) Wright-Giemsa-stained cytospins of BM cells from leukemia mice receiving free BMS, HDL-AuNPs-BMS or vehicle (original magnification × 400). Graph is the quantification of post-mitotic cells. (h,j) qPCR analysis of *DNMT1* and *p15*^*INK4B*^ expression from BM cells of leukemia mice receiving free BMS, HDL-AuNPs-BMS or vehicle. (i) Dotblot of genomic DNA from BM cells of leukemia mice receiving free BMS, HDL-AuNPs-BMS or vehicle. In all experiments, n = 3 mice/group, data are mean ± SD; \**P* < 0.05, \*\**P* < 0.01; BMS represents BMS309403; N-BMS represents HDL-AuNPs-BMS. Note: Emp, F-BMS, N-BMS represents empty vehicle, free BMS, HDL-AuNPs-BMS, respectively.

## 4. CONCLUSION

In this study, we have successfully prepared the first HDL-AuNPs loaded with small molecule inhibitor. The *in vitro* and *in vivo* results herein for the first time demonstrate that HDL-AuNPs are an effective vehicle for delivering small molecule inhibitors (e.g., BMS) to cancer cells naturally targeted by HDLs. Our data clearly demonstrate proof-of-concept of the function of the HDL-AuNPs-BMS to suppress leukemia growth through the deployment of BMS to restore normal DNA methylation profile, thus offering a paradigm-shift for treating leukemia. Given that SRB1 is silenced in the majority of normal human tissues,^17^ but consistently overexpressed by most tumor cells,^49^ our encouraging findings provide the basis to further exploit HDL-AuNPs-BMS in SRB1-mediated delivery to different types of cancer characterized by FABP4 upregulation.

## Acknowledgements

This work was supported partially by The Hormel Institute Foundation and National Cancer Institute (Bethesda, MD) grants R01CA149623, R21CA155915 and R03CA186176. The electron micrographs were collected using a Tecnai TF30 transmission electron microscopy (TEM) maintained by the Characterization Facility, University of Minnesota.

## Author Contributions

S.J.L conceived ideas, designed the experiments and oversaw the entire research project; N.S., F.Y., and J.X.P performed experiments; Z.G. and C.L.H characterized the HDL-AuNPs by transmission electron microscopy; A.A. and M.R.L provided the patient samples and critically reviewed the manuscript. N.S., F.Y., C.L.H and S.J.L analyzed data and wrote the paper; N.S. and F.Y performed the biostatistical analysis.

